# Effects of diversifying the planting period of maize on herbivorous and their natural enemies

**DOI:** 10.1101/2022.01.03.474813

**Authors:** Gemma Clemente-Orta, Hugo Alejandro Alvarez, Filipe Madeira, Ramon Albajes

**Affiliations:** Department of Crop and Forest Sciences, AGROTECNIO Center, University of Lleida, Rovira Roure 191, 25198 Lleida, Spain; Department of Zoology, University of Granada, Granada, Spain; Environmental and Ecosystem Management Area, MORE CoLab - Mountains of Research Collaborative Laboratory, Av. Cidade de Léon 596, 5300-358 Bragança, Portugal

**Keywords:** crop rotation programs, integrated pest management, crop phenology and planting periods

## Abstract

Knowledge of the specific insect densities during crop development is necessary to perform appropriate measures for the control of insect pests and to minimize yield losses. In a previous study, both spatial and temporal approaches were adopted to analyse the influence of landscape structure and field variables on herbivore and predatory insects on maize. Both types of variables influenced insect abundance, but the highest effect was found with maize phenology. Given that the field planting date could modulate the influence produced by the structure of the landscape on herbivores and predatory insects, analyses of population dynamics must be performed at both the local and landscape levels. The anterior prompted us to study these aspects in the two common planting periods (early and late) in northern Spain. The present study tests the hypothesis that the period of maize planting could have a higher effect than phenology or interannual variation on the abundance of natural enemies and herbivores on maize. Our results showed that only the abundances of ‘other herbivore thrips’ and Syrphidae were significantly different between the two planting periods. Moreover, we found significant effects of planting period when we performed yearly analysis in 2015 for Coccinellidae and Chrysopidae and in 2016 and 2017 for *Aeolothrips* sp. Most of the taxa had abundance peaks in earlier growth stages, which are related to pollination (before or during), while only *Stethorus punctillum* and Syrphidae increased later in the season. Furthermore, *Frankliniella occidentalis*, aphids, Syrphidae and Coccinellidae registered higher abundances in fields sown in the late planting period than in the rest of the insect species. The results of the present study highlight the effects of sowing dates on insect dynamics in maize.

**Highlights:**

- Herbivorous and natural enemy abundance peaks in earlier growth stages.
- Abundance of thrip and Syrphidae were different between the two planting periods.
- Abundance of Coccinellidae, Chrysopidae and *Aeolothrips* sp. was different when performed yearly analysis.

## Introduction

Maize is the most common arable crop in summer in the Ebro Basin (Northeast Spain). Farmers traditionally plant maize rather early in spring, after the winter fallow, to increase its yield and ensure a correct low grain humidity at harvesting (Maresme et al., 2019). In recent years, however, due to the increasing irrigated surface of agricultural land, the longer growing season caused by the warmer conditions in the area, and the decreasing revenue received by cereal growers, the rotation based on winter and summer cereals is intensifying. As a consequence, maize is increasingly planted after the cereal harvest during the spring in comparison with the previous common habit of planting after the winter fallow at the beginning of spring. Whereas maize can be planted from the end of March to early July, it is harvested for grain or forage from the end of September to November or even in early December in mild autumns.

Therefore, in the same area, fields with different phenologies coexist side by side so that most herbivores and their natural enemies (NEs) can move in the landscape to select the preferred maize plant age for feeding and reproduction. Many studies have determined optimal planting dates and maize cultivar cycles to maximize yields (Tsimba et al., 2013; Hall et al., 2016; Kucharik et al., 2006), but studies have rarely addressed the influence of maize planting dates on insect pests and viral diseases. Knowledge of how planting date may affect the composition and population dynamics of maize herbivores and their NEs may contribute to understanding and predicting the impact of climate change and the alteration of agricultural practices on maize insect pests and viruses vectored by insects.

Herbivorous insects affect crop yield due to both their plant feeding and virus transmission capacity. Among herbivores feeding on maize, borers (Lepidoptera) and homopterans (Hemiptera) are the most damaging pests in the region (MAAM., 2015). Maize borers in the region belong to two species, *Sesamia nonagrioides* Lef. (Lepidoptera: Noctuidae) and *Ostrinia nubilalis* (Hbn) (Lep.: Pyralidae). Chemicals are rarely applied to control these species because of their poor efficacy due to the endophytic habits of borer larvae, the difficulty of application due to crop height, and their strong impact on natural enemies (NEs) (Asin and Pons, 1999; Albajes et al., 2003; Ardanuy et al., 2018). Insect-resistant cultivars, particularly genetically modified (GM) maize, have been the most successful control method in recent years (Bt maize, Hellmich, et al., 2007).

Other relevant damaging insects on maize in the area include Homopterans, both Auchenorrhyncha and Sternorrhyncha (Albajes et al., 2013). Among Auchenorrhyncha, the planthopper *Laodelphax striatellus* (Fallén) (Hemiptera: Delphacidae) is harmful to maize mainly due to its capacity to transmit maize rough dwarf virus (MRDV) (Achón et al., 2013, 2015), and the leafhopper *Zyginidia scutellaris* Herrich-Schäffer (Hemiptera: Cicadellidae) causes plant vigour reduction due to its feeding activity on the phloem (mostly in early plant growth stages, Pons and Albajes, 2002). The main group of Sternorrhyncha affecting maize yield in the area includes several species of aphids (Hemiptera: Aphididae), the phenology of which depends on the species, but in general, they are more abundant before anthesis (Pons et al., 1994); their main damage comes from their high capacity to transmit two common maize viruses, maize dwarf mosaic virus (MDMV) and sugarcane mosaic virus (SCMV) (Achon and Sobrepere, 2001), the two main potyviruses in this area. Although soil insecticides may cause significant homopteran suppression (Pons and Albajes, 2002), they are only partially effective in preventing the incidence of aphid-transmitted viruses due to their nonpermanent transmission (Asin and Pons, 1999). In contrast, insecticide treatments on maize showed negative effects on NE abundance (Asin and Pons, 1999; Albajes et al., 2003).

In a recent study, Clemente et al. (2020a) investigated the influence of the surrounding landscape and crop fields on the abundance of maize herbivore insects and their NE. They showed that the variable with the highest effect on insect abundance was the maize growth stage. Specifically, in spring, the abundance of the predators *Orius* sp., *Propylea quatuordecimpunctata*, and Chrysopidae was positively related to crop phenology, while it was negatively related to *Stethorus* spp. and Syrphidae. Conversely, in summer, positive relationships were found only for *Stethorus* spp., whereas negative relationships were found for *P. quatuordecimpunctata* and *Aeolothrips* sp. In addition, phytophagous Thripidae (except *Frankliniella* sp.), *Empoasca vitis*, aphids, phytophagous Thripidae, *Z. scutellaris*, and *L. striatellus* showed negative relationships in both spring and summer. Moreover, it was found that early and late maize planting dates have important effects on the incidence of SCMV, MDMV (Clemente-Orta et al., 2020b), and MRDV (Clemente-Orta, et al., 2021).

The small amount of available information regarding the influence of maize planting date on the seasonal abundance of herbivores prompted us to study these aspects in two current common planting periods in northeastern Spain. In the present study, we tested the hypothesis that the planting period of maize could have a greater effect on the abundance of herbivores and their NEs than maize phenology or interannual variation. To check this hypothesis, we sampled 52 maize fields planted on different dates for 3 consecutive years.

## Materials and Methods

### Study area and cultivation practices

This study was carried out in 2015, 2016 and 2017 in the Ebro Basin (41°48′12.20″N, 0°32′45.77″E; 120–346 m altitude; 200–400 mm rainfall, Tmin: 8°-24 °C and Tmax: 18°-38 °C) (Fig. 1a). In the region, the most traditional rotation included winter and summer cereals and alfalfa. Pest management in cereals includes pre- and postemergence herbicide applications, treatment of seeds of winter cereals with fungicides, and treatment of maize with both insecticides and fungicides. Under these conditions, a total of 52 maize fields were selected to have a wide variety of planting dates in such a way that several growth stages coexisted throughout the season in the area (Fig. 1b). The sizes of the fields varied between 0.9 and 13.68 ha, and fields were located at least 2 km apart from each other. The growth stage interval of maize plants was recorded with the Ritchie et al. (1992) nomenclature. For analysis, fields were grouped into two groups: the early planting period included those sown from March until the end of April, whereas late-planted fields were those planted during May or June (Fig. 1c).

**Figure 1.**
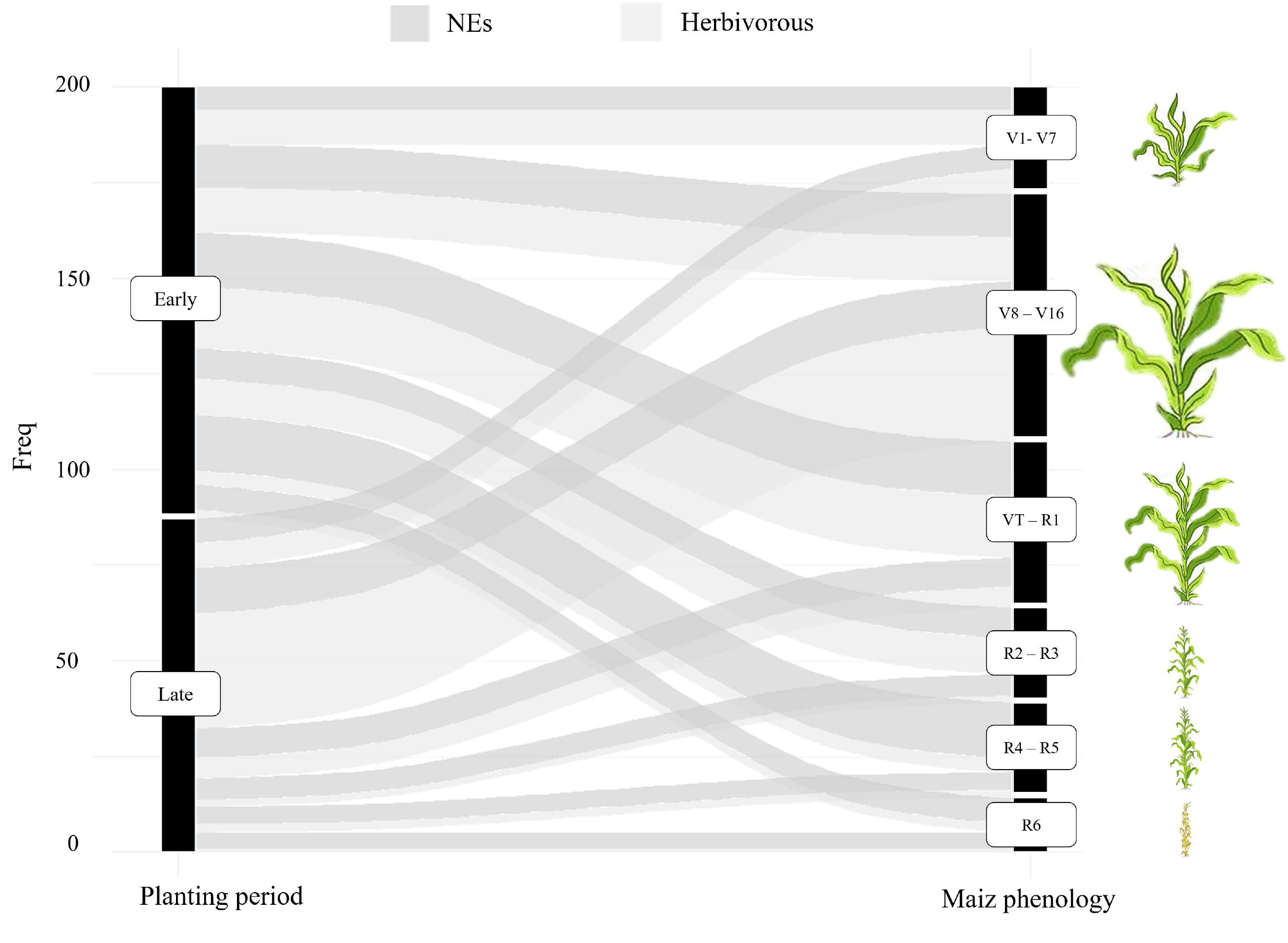
Maize growing areas in Ebro valley, northern Spain. (a) Study area, (b) maize fields sampled, and (c) the two common planting periods in the area (early and late) with maize phenology.

### Sampling herbivores and natural enemies

The abundance of insects in maize fields was estimated using yellow sticky traps (30 ×25 cm, Pherocon Trece, Adair, Oklahoma). Sampling was carried out in spring and summer in 2015, 2016, and 2017. Sampling was performed each year within the following date (sampling date SD) intervals: SD1: 15 May-15 June; SD2: 22 June-10 July; SD3: 10-22 August; SD4: 31 August-20 September. Crop growth stages were grouped into 6 intervals: (1) V1-V7; (2) V8-V16; (3) VT-R1; (4) R2-R3; (5) R4-R5; and (6) R6.

Traps were left active for 1 week. In each field, 3 traps were placed at crop canopy height along a transect perpendicular to the nearest edge (stacked on stakes approximately 30 m apart), and traps were separated from each other by 15 m (Albajes et al., 2013). After that, traps were collected and stored at 6-8 ºC until insect identification. Insects were identified at different taxonomy levels: family, genus or species.

### Statistical analysis

We fitted generalized linear mixed models (GLMMs) to analyse insect population dynamics (as repeated measures), for which we used weekly mean numbers of the three traps placed in each field as the dependent variable in all models using the negative binomial tendency. In the first analysis aimed to compare the effect of planting date on insect abundance, we included the planting periods (early and late as defined above), maize phenology, and their interaction as fixed effects. In this analysis, site identification (ID) with and without year were included as random effects to assess the effect for all years (overall effect) and year by year, respectively. In a second type of analysis aimed to identify the interannual variation (temporal effect), we included year as a fixed effect and maize phenology and site ID as random effects in a second analysis. Further differences between the groups in each model were tested using contrasting *post hoc* tests.

All analyses were computed in R software v 3.6.2 (R Developmental Core Team, 2019). For each model, we tested significant differences using the ANOVA function of the car package. GLMMs and *post hoc* tests were computed using the lme4 and multcomp packages, respectively.

## Results

A total of 316,564 insects were trapped on 585 yellow sticky traps placed in the 52 maize fields during the three years of study. Herbivorous insects identified in traps included *Frankliniella occidentalis*, ‘other herbivore thrips’ (others different from *Frankliniella* sp.), *Zyginidia scutellaris, Laodelphax striatellus, Empoasca* sp., and aphids. The NEs identified were *Orius* sp., *Nabis* sp., Miridae, *Stethorus punctillum, Chrysoperla* sp., Syrphidae, *Aeolothrips* sp. and Coccinellidae. The total numbers of insects caught in traps per year were 201,775 in 2016 (*n* = 23 fields), 75,250 in 2017 (*n* = 23) and 39,539 in 2015 (*n* = 6).

In contrast to that initially hypothesized, the planting period had no significant effects on insect catch numbers for any of the taxa recorded except for the group of ‘other herbivore thrips’ (Table 1), in which fields planted earlier showed higher numbers. However, when this factor interacted with phenology, 5 taxa exhibited significant results (Tables 1 and 2). When the influence of the planting period on insect catches was analysed within each of the three years, only in 3 of the 36 possible cases (12 insect taxa per 3 years) were there significant abundance differences between planting periods: in 2015 for Coccinellidae (χ^2^ = 6.76; df = 1; *p* = 0.0002) and Chrysopidae (χ^2^ = 7.94; df = 1; *p* = 0.004) and for *Aeolothrips* sp. in both 2016 (χ^2^ = 5.02; df = 1; *p* = 0.02) and 2017 (χ^2^ = 11.7; df = 1; *p* = 0.006), with higher values in the earlier-planted fields in all three cases.

**Table 1.** Results of the generalized linear mixed models of herbivore abundance according to maize phenology and planting period (early and late) and their interaction for all years.

| Herbivore | Variable | Chisq | Df | p value |
| --- | --- | --- | --- | --- |
| *F. occidentalis* | Planting | 1.437 | 1 | 0.23 |
|  | Phenology | 147.29 | 5 | **<0.001** |
|  | Planting:Phenology | 210.84 | 5 | **<0.001** |
| “Other herbivore thrips” | Planting | 12.62 | 1 | **<0.001** |
|  | Phenology | 494.35 | 5 | **<0.001** |
| *Z. scutellaris* | Planting | 0.77 | 1 | 0.38 |
|  | Phenology | 160.92 | 5 | **<0.001** |
| *Empoasca* sp. | Planting | 0.201 | 1 | 0.65 |
|  | Phenology | 18.08 | 5 | **0.002** |
|  | Planting:Phenology | 11.98 | 5 | **0.03** |
| Aphids | Planting | 3.31 | 1 | 0.06 |
|  | Phenology | 564.93 | 5 | **<0.001** |
| *L. striatellus* | Planting | 0.42 | 1 | 0.51 |
|  | Phenology | 66.36 | 5 | **<0.001** |
|  | Planting:Phenology | 15.3 | 5 | **0.009** |

**Table 2.**
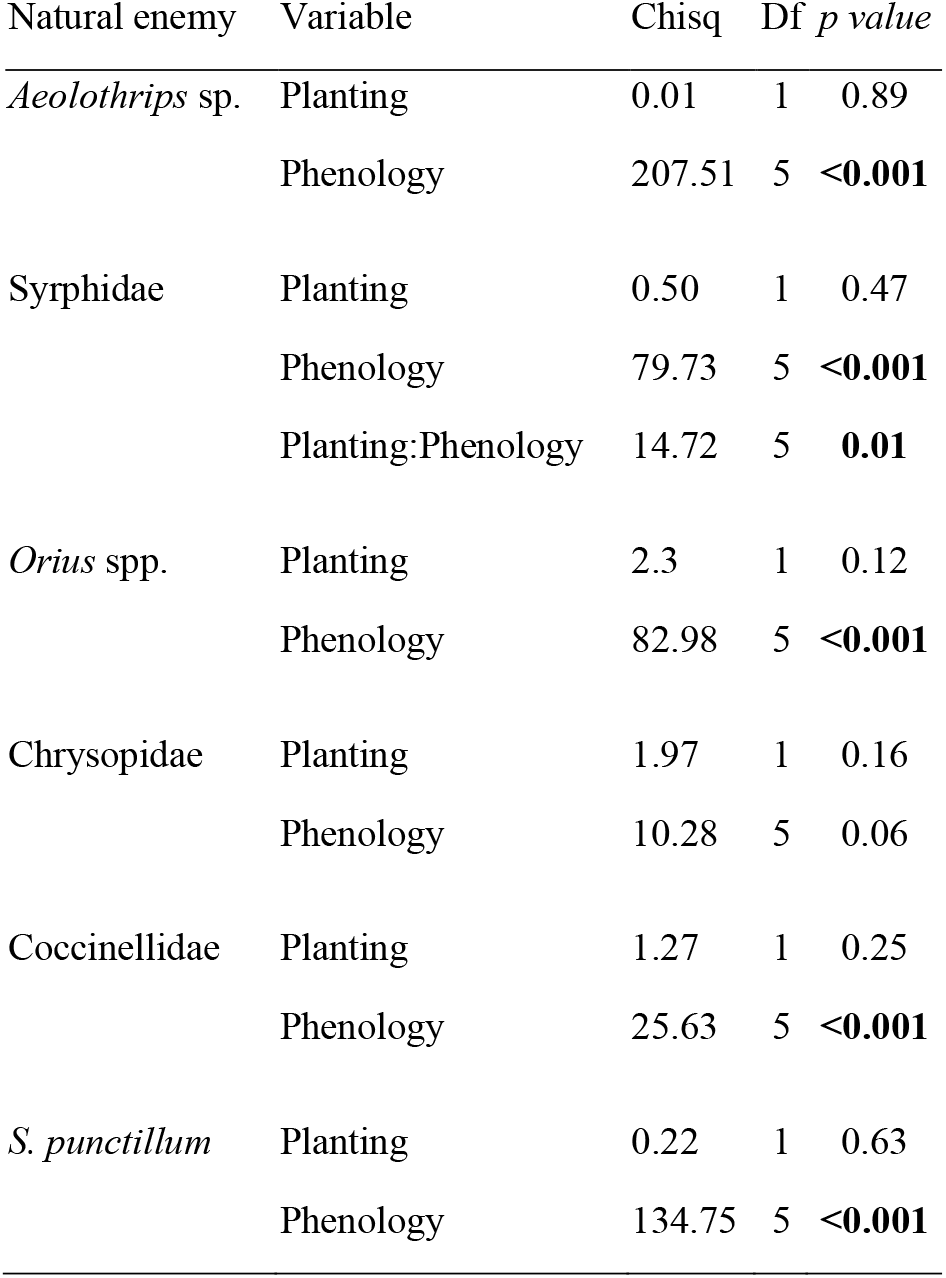
Results of the generalized linear mixed models of NE abundance according to maize phenology and planting period (early and late) and their interaction for all years.

| Natural enemy | Variable | Chisq | Df | *p value* |
| --- | --- | --- | --- | --- |
| *Aeolothrips* sp. | Planting | 0.01 | 1 | 0.89 |
|  | Phenology | 207.51 | 5 | **<0.001** |
| Syrphidae | Planting | 0.50 | 1 | 0.47 |
|  | Phenology | 79.73 | 5 | **<0.001** |
|  | Planting:Phenology | 14.72 | 5 | **0.01** |
| *Orius* spp. | Planting | 2.3 | 1 | 0.12 |
|  | Phenology | 82.98 | 5 | **<0.001** |
| Chrysopidae | Planting | 1.97 | 1 | 0.16 |
|  | Phenology | 10.28 | 5 | 0.06 |
| Coccinellidae | Planting | 1.27 | 1 | 0.25 |
|  | Phenology | 25.63 | 5 | **<0.001** |
| *S. punctillum* | Planting | 0.22 | 1 | 0.63 |
|  | Phenology | 134.75 | 5 | **<0.001** |

Maize phenology (indicated by the growth stage intervals at the sampling week) had significant effects on the abundance of most NEs (except Chrysopidae) (Table 2) and all herbivores (Table 1) by itself or interacting with planting date. In particular, the number of insects caught on traps varied with crop growth stage, with clear peaks in certain growth stages (Fig. 2). Most of the herbivores and NEs peaked before or at pollination in both earlier- or later-planted fields. The herbivore groups of ‘other herbivore thrips’ together with aphids and Syrphidae showed their highest abundances in the earliest growth stage (V1-V7). whereas the genus *Empoasca* sp. peaked in the first or second growth stage intervals in fields planted earlier or later, respectively. The herbivore thrips *F. occidentalis* showed only a high peak in fields planted later, which occurred in the second growth stage interval, whereas the rest of the catches were low throughout the entire season independent of the planting period. The predatory thrips *Aeolothrips* sp. peaked at the second growth stage interval in both early- and late-planted fields. The rest of the herbivores and NEs peaked around the pollination growth stage and independent of the planting date; only Chrysopidae, especially *S. punctillum*, peaked after pollination and reached significantly higher numbers in R4-R5.

**Figure 2.**
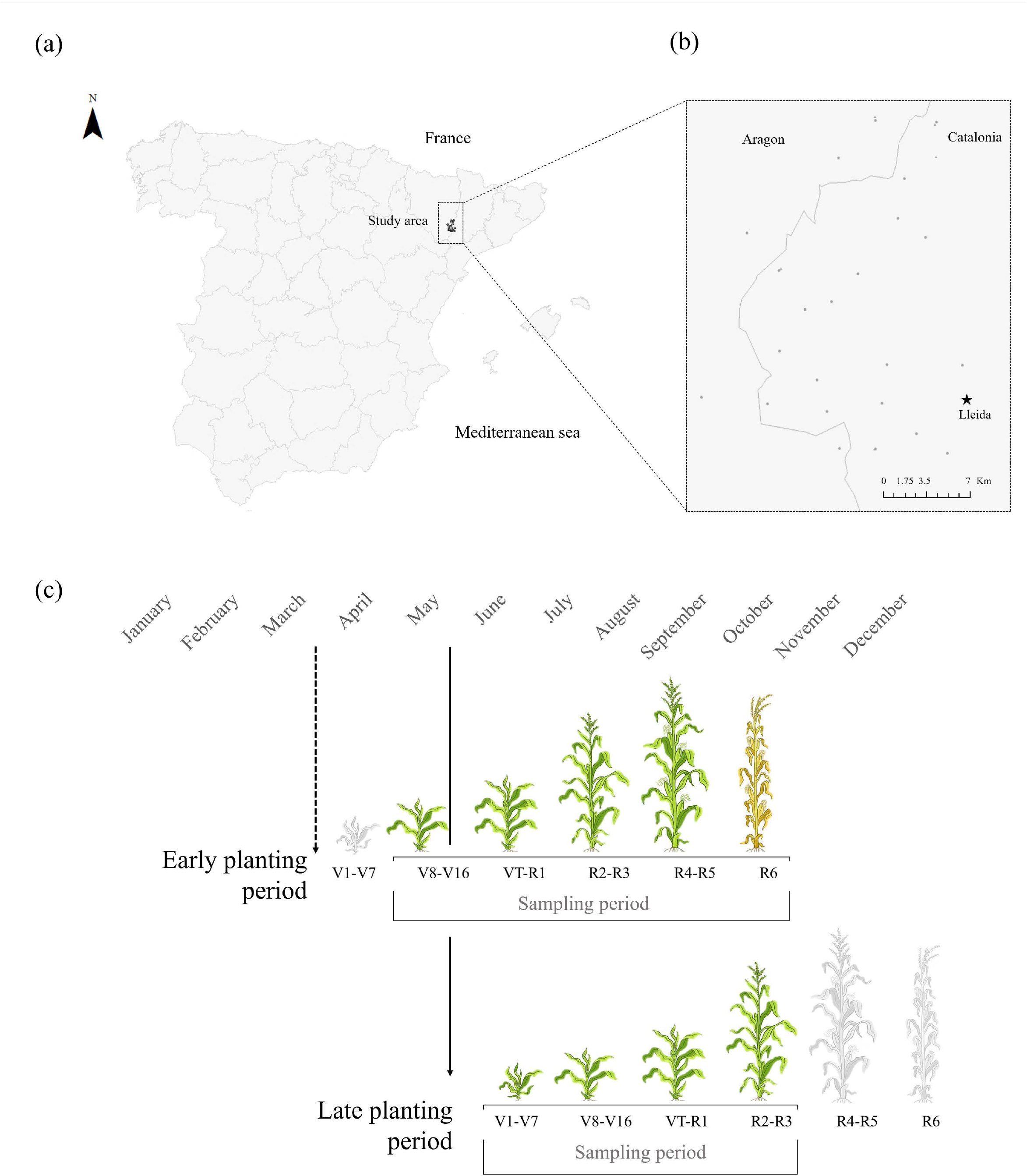
Insect abundance according to maize phenology for all years in each planting period (early and late) with the mean and standard deviation. The letters show significant differences among phenologies. Asterisks indicate that there are differences between planting periods. Chrysopidae show non-significant (ns) results.

The second type of analysis was performed to observe interannual variation in insect abundance along the season by year. Overall, the dynamics of insects during the season were quite consistent in the three years for most of the taxa recorded. Moreover, insect prevalence was quite similar in the three years for both herbivores and NEs (Supplementary material Fig. 1 and Fig. 2). Among herbivores, the highest numbers were registered for *F. occidentalis*, followed by *Z. scutellaris*; for NEs, *Orius* sp. was the most abundant taxon, followed by *Aeolothrips* sp. and *S. punctillum*. The analysis of interannual variation also showed that the abundance of insects collected in early *vs*. late planting periods followed a rather similar pattern, although it varied significantly in the cases of 5 herbivores, *F. occidentallis, Z. scutellaris, L. striatellus, E. vitis* and, Aphididae (Fig. 3), and 3 NEs, *Orius* sp., *Aeolothrips* sp., and Coccinellidae (Fig. 4).

**Figure 3.**
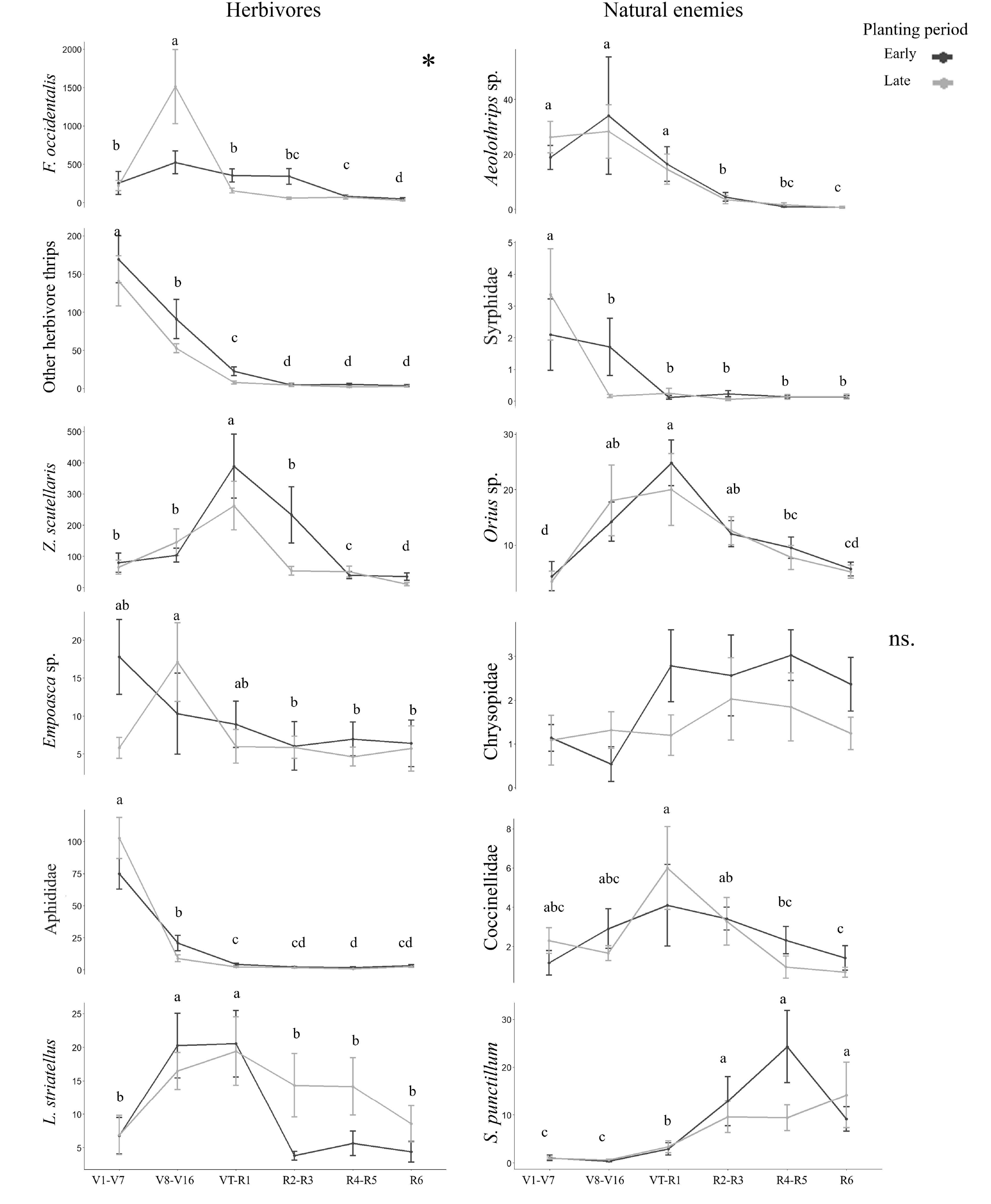
Herbivore abundance according to maize phenology each year in each planting period (dashed line = early; continuous line = late). The species that had significant differences in the analyses are shown.

**Figure 4.**
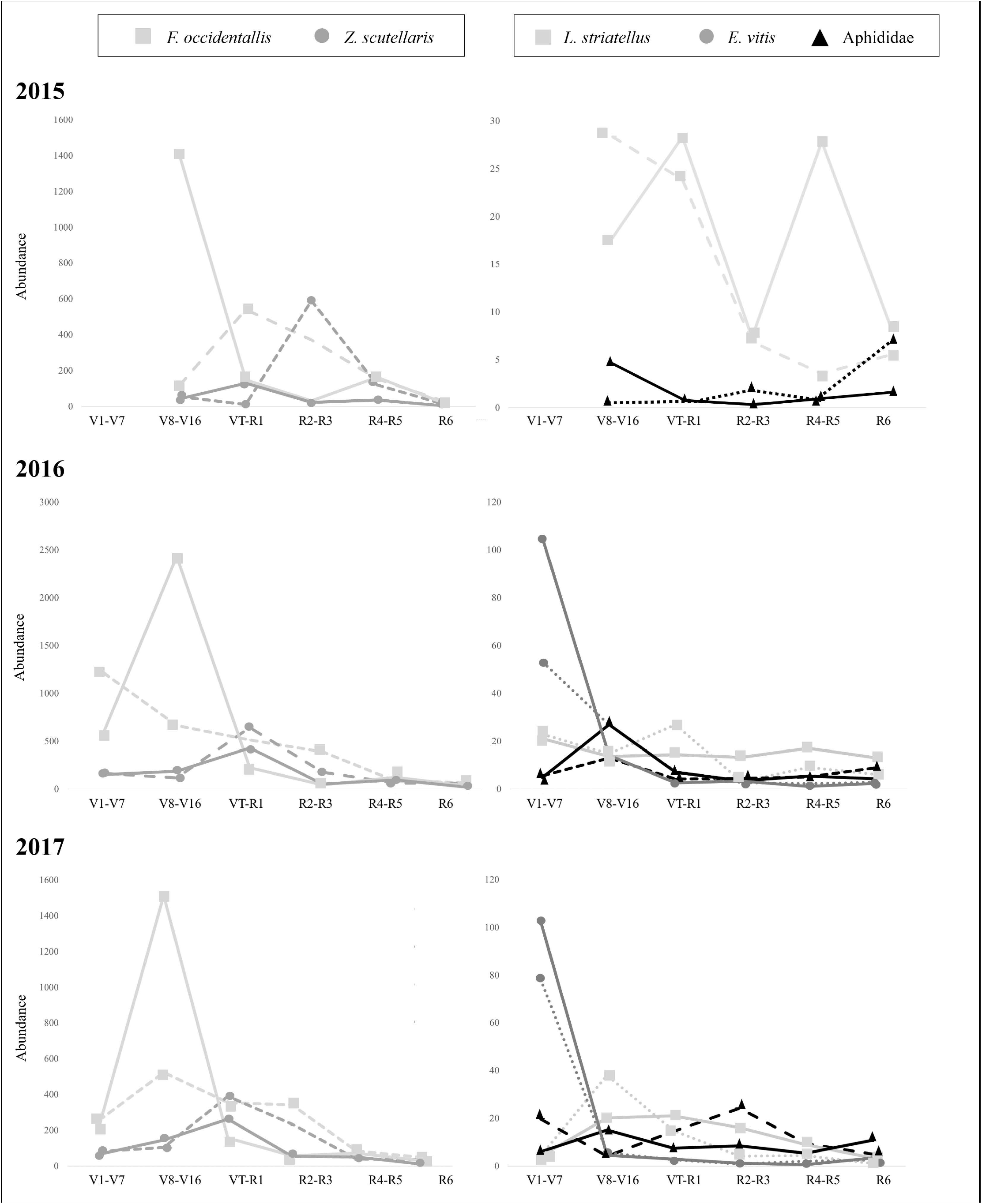
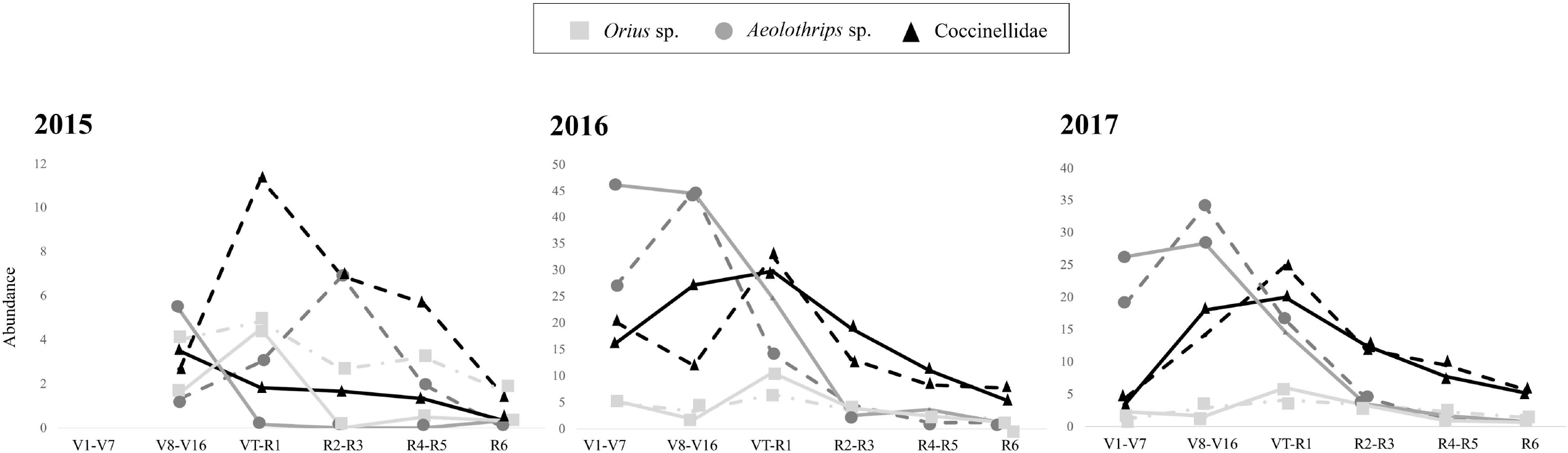
Natural enemy abundance according to maize phenology each year in each planting period (early and late). Only the species that had significant differences in the analyses are shown.

## Discussion

The diversity and abundance of maize insect taxa caught on yellow sticky traps were within the ranges reported by authors in previous studies conducted in regions with yellow sticky traps (Albajes et al., 2009, 2013) or by visual on-plant sampling over the last 15 years (Albajes et al., 2003; Pons et al., 2005; Poza et al., 2005). Among herbivores, thrips and hemipterans were predominant, whereas among predators, the generalist *Orius* sp., predatory thrips, and the mite-feeding specialist *S. punctillum* were the most abundant taxa recorded.

Although we initially expected that planting period would have a strong effect on total insect abundance on the crop, when we analysed this factor, we did not find remarkable effects in the different models for most of the taxa recorded. This result could indicate that damage caused by herbivores on the crop is not strongly affected by the maize planting period, and an important conclusion involves the recent frequent variation in planting dates of maize in the study area. However, differences in insect abundance according to the planting period were found at particular growth stages, a remarkable feature in terms of crop damage potential caused on maize, as crop plants have different susceptibilities and yield responses to herbivores and disease vectors during their development (Liliane and Charles, 2020). For example, *F. occidentalis* was approximately three times more prevalent in fields planted later than in early plantings at V8-V16, when plants were more susceptible to thrips feeding. This differential susceptibility of maize plants to herbivore insect activity at different growth stages has been particularly observed for vectors of maize potyviruses (Achon and Sobrepere, 2001; Clemente-Orta et al., 2020a) and maize rough dwarf disease (MRDV) (Achon et al., 2013, 2015; Clemente-Orta et al., 2020b). In the first case, aphids that are able to transmit maize potyviruses were not found to be influenced in their abundance by the planting period at any growth stage in the present study. In the case of MRDV, its vector, *L. striatellus*, did show significantly higher catches in late-planted fields but only at late growth stages, when virus transmission is already complete (Achon et al., 2013, 2015) and the number of vectors rather irrelevant when considering virus incidence.

Maize phenology was the most influential factor on insect abundance. The numbers of most insects peaked before or during pollination, except in the case of *S. punctillum*, which peaked at R4-R5, likely soon after the spider mite population peak. Little is known in general about the suitability of maize plants for insect herbivores during their development in the field. Most information on this issue classically refers to the role of hydroxamic acid (mostly DIMBOA) contents in maize plants on herbivores, mainly studied for maize borers (Gutierrez and Castañera, 1986; Barry et al., 1994) and aphids (Thackray et al., 1990). However, as contents in the DIMBOA plant and related compounds decrease with plant age (Cambier et al., 2000), it does not seem that its role in shaping the abundance of insect herbivores is crucial, and other causes of insect decline after maize pollination should be investigated to understand the herbivore insect preference for some maize growth stages. Most NEs recorded in this study peaked soon after herbivore prey did, as in the case of *Aeolothrips* sp. and *F. occidentalis*, ‘other herbivore thrips’, and the case of *Orius* sp. and hemipterans, a phenomenon already noted in maize crops by Albajes et al. (2011) and Ardanuy et al. (2018).

The composition and configuration of the landscape could modulate the influence of field planting date on herbivores and NE insects (Clemente-Orta et al., 2020) as a consequence of insect movement among habitats, resulting in spatial or temporal migrations (Tscharntke et al., 2012). The combination of many trophic-level interactions, the landscape structure, the management of crop fields and the constant changes in agricultural policy make it difficult to fully understand and predict the changing patterns of insect abundance in particular agricultural habitats.

We could conclude that a number of potential consequences derived from varying maize planting periods can be expected. However, the pressure of insect pests on maize and the total number of NEs did not change substantially according to the planting period, whereas most consequences will likely come from the different susceptibilities of maize plants of different ages to insects and insect-virus vectors. Most insect populations studied in this work peaked before or during maize plant pollination, as Albajes et al. (2009) reported in previous studies. In this period, high differences in plant phenology may lead to yield reduction. In parallel, although the number of generalist predators was not greatly influenced by the differential amount of prey (herbivores) according to the planting period, the specialist NEs may be influenced by it. Finally, considerations about landscape structure, as well as field and crop management practices, may also modulate the impact of maize planting date on insect dynamics; therefore, further studies on the influence of planting date on insect herbivores and their NEs should include both factors.

## Acknowledgements

This research was funded by the Spanish Ministry of Economy, Industry and Competitiveness project AGL2014-53970-C2-1-R. and AGL2017-84127-R. G. Clemente-Orta was funded by the grant BES-2015-072378 from the Ministry of Science, Innovation and Universities. Technicians for the agricultural cooperatives are acknowledged for providing information on the management, and the landowners, for allowing us to access to their fields. We also thank two anonymous reviewers whose comments have greatly improved this manuscript.

## Author contribution

R.A., F. M. and G.C.O. contributed to study design and collected the data. R.A., G.C.O. and H.A.A performed the data analysis and, wrote the manuscript. All authors read, revised and approved the final version of the manuscript.

## Notes

### Competing Interest Statement

The authors have declared no competing interest.

